# Process-explicit models reveal pathway to extinction for woolly mammoth using pattern-oriented validation

**DOI:** 10.1101/2021.02.17.431706

**Authors:** Damien A. Fordham, Stuart C. Brown, H. Reşit Akçakaya, Barry W. Brook, Sean Haythorne, Andrea Manica, Kevin T. Shoemaker, Jeremy J. Austin, Benjamin Blonder, Julia Pilowsky, Carsten Rahbek, David Nogues-Bravo

**Author notes:** **Corresponding author:** Damien Fordham, University of Adelaide, Adelaide, South Australia, Australia, 5005 (**email:**; <u>ph:</u>; <u>fax:</u>). Statement of authorship:* D.A.F, D.N.B. and C.R. conceived the idea for the paper. D.A.F. and S.C.B. did the analysis with support from H.R.A, B.W.B, S.H., A.M., K.T.S, J.J.A., J.P. and B.B. All authors contributed to writing the manuscript. Data accessibility statement:* Data supporting the results are archived on Figshare (http://doi.org/10.25909/5f22592242ca2) and the R-code for the process-explicit model is archived on Zenodo (https://doi.org/10.5281/zenodo.5567859).

## Abstract

Pathways to extinction start long before the death of the last individual. However, causes of early-stage population declines and the susceptibility of small residual populations to extirpation are typically studied in isolation. Using validated process-explicit models, we disentangle the ecological mechanisms and threats that were integral in the initial decline and later extinction of the woolly mammoth. We show that reconciling ancient DNA data on woolly mammoth population decline with fossil evidence of location and timing of extinction requires process-explicit models with specific demographic and niche constraints, and a constrained synergy of climatic change and human impacts. Validated models needed humans to hasten climate-driven population declines by many millennia, and to allow woolly mammoths to persist in mainland Arctic refugia until the mid-Holocene. Our results show that the role of humans in the extinction dynamics of woolly mammoth began well before the Holocene, exerting lasting effects on the spatial pattern and timing of its range-wide extinction.

## Introduction

Despite knowledge from the early 19^th^ century (Cuvier 1807) that species go extinct, ecological mechanisms that underpin extinctions remain poorly resolved (Beissinger 2000; Channell & Lomolino 2000; Traill *et al*. 2007). This is because pathways to extinction can begin long before the extinction event, resulting from driver-state relationships that are difficult to detect and disentangle (Soulè 1983; Caughley 1994). In contrast, the demographic, ecological and genetic processes that make small populations susceptible to eventual extinction are better established (Lande 1993; Frankham 2005). Here we develop a process-explicit modelling framework that integrates the declining and small population paradigms, central to ecology and conservation biology (Caughley 1994), using pattern-oriented validation (Grimm *et al*. 2005). We apply it to the range collapse and extinction of the woolly mammoth (*Mammuthus primigenius*) to develop a more holistic understanding of spatiotemporal extinction dynamics. The woolly mammoth was one of many large mammals that went extinct during the Pleistocene-Holocene transition (Barnosky *et al*. 2004; Cooper *et al*. 2015).

Global warming following the Last Glacial Maximum, which extended from the late Pleistocene to the early Holocene, resulted in regional temperature increases of 4 to > 10 °C (Clark *et al*. 2012). During this deglaciation phase, many megafaunal species (terrestrial taxa > 45 kg) became extinct (Barnosky *et al*. 2004) and many others suffered regional extirpations (Cooper *et al*. 2015). This biotic simplification radically changed the structure and function of ecosystems (Gill *et al*. 2009; Doughty *et al*. 2016). At the same time, Palaeolithic human populations were spreading and becoming more ubiquitous, facilitated by increases in primary productivity associated with climatic change (Eriksson *et al*. 2012; Timmermann & Friedrich 2016). What has remained fiercely contested is the relative role of human hunting and climate change, or a synergy of these impacts, on the fate of the megafauna (Barnosky *et al*. 2004; Stuart *et al*. 2004; Stuart 2005; Nogués-Bravo *et al*. 2008; Lorenzen *et al*. 2011; MacDonald *et al*. 2012; Cooper *et al*. 2015). Major barriers to a resolution have included a sparse and uncertain fossil record (Haile *et al*. 2009), a lack of high-resolution spatiotemporal projections of climatic change and human abundances (Fordham *et al*. 2018), reliance on correlative rather than process-based approaches to infer drivers of extinction from ecological and molecular data (Fordham *et al*. 2020), and a focus on extinctions of populations once at critically small thresholds (Palkopoulou *et al*. 2015; Graham *et al*. 2016; Rogers & Slatkin 2017), rather than the causes of smallness itself (Caughley 1994).

Recent developments in macroecological modelling are enabling the drivers and processes of megafauna extinctions to be unravelled from the point of initial population decline to the final extinction event, using a wide body of evidence from paleo-archives (Fordham *et al*. 2020). These novel approaches, which simulate the dynamics of an ecological system as explicit driver-state relationships (Rangel *et al*. 2018), offer fresh opportunities to disentangle the mechanisms responsible for ecological responses to climate- and human-driven changes in species distributions and abundances across space and time (Fordham *et al*. 2016a). Unlike correlative approaches, fossil and molecular signatures of past demographic change are used as independent, objective targets for directly evaluating a model’s structural adequacy and parameterization (Grimm *et al*. 2005). This new pattern-oriented approach to refining and validating process-explicit models of species’ range dynamics allows relevant global change drivers and ecological processes to be simulated and tested, revealing the most likely chains of causality that lead to extinction (Fordham *et al*. 2021).

The iconic woolly mammoth was present on earth for more than half a million years (Bevan *et al*. 2017) before going extinct in the mid-Holocene (Stuart *et al*. 2004). During this time, woolly mammoths co-existed with Neanderthals (*Homo neanderthalensis*) and modern humans (*H. sapiens*) for many millennia, and were exploited for meat, skins, bones and ivory (Stuart *et al*. 2004; Stuart 2005; Nogués-Bravo *et al*. 2008). As the earth warmed rapidly during the last deglaciation (approximately 19 to 11 ka (thousand years ago) BP), boreal forests spread throughout Eurasia, replacing tundra grassland and forbs (Binney *et al*. 2017), the preferred habitat for woolly mammoths.

Previous process-explicit examinations of the extinction dynamics of woolly mammoth focused largely on threats to persistence for populations at already critical thresholds (Palkopoulou *et al*. 2015; Graham *et al*. 2016; Rogers & Slatkin 2017), concluding that genomic meltdown through inbreeding caused its extinction (Palkopoulou *et al*. 2015; Rogers & Slatkin 2017). Where long-lasting roles of climate and humans have been considered, their function in setting the location and timing of woolly mammoth extinction (and extinctions of other large mammals) during the Pleistocene-Holocene transition has been inferred from snapshots (points in time ≥ 12,000 years apart) of projected historical range movement (Nogués-Bravo *et al*. 2008; Lorenzen *et al*. 2011) and analysis of time-binned fossil and archaeological data (Stuart *et al*. 2004; MacDonald *et al*. 2012), using correlative (not process-explicit) modelling methods. Consequently, the spatiotemporal dynamics of extinction forces, and resultant long-term patterns in population and range collapse, remain unclear.

To unravel the determinants of early-stage population declines, and subsequent range collapse and extinction of woolly mammoth in Eurasia, we built process-explicit spatially dynamic macroecological models that simulate how ecological requirements and demographic processes interact with climate change and human pressures to affect the geographical range, population dynamics and the range-wide extirpation pattern of woolly mammoths from 21 ka BP (kilo annum Before Present). We explored more than 90,000 scenarios, deriving best estimates of model parameters with pattern-oriented validation methods. As validation targets, we used spatiotemporal inferences of local extinctions and demographic change from hundreds of radiocarbon dated fossils and ancient DNA sequences. These validation targets identified models with the combinations of ecological processes (niche and demographic constraints for movement and extinction) and rates of global change (climate-change, human-impact and their interaction) that best reconcile with ancient DNA data on the timing and rate of population decline, along with fossil evidence of the extirpation pattern, and location and timing of eventual species extinction. In doing so, our new macroecological modelling approach was able to directly disentangle in space and time the processes and threats crucial to the initial population decline and later extinction of the woolly mammoth, revealing that its pathway to extinction started long before the final extinction event.

## Materials and Methods

The process-explicit macroecological model of climate-human-woolly mammoth interactions we built was designed to reconstruct the pattern of range collapse, population decline and extinction of woolly mammoth in Eurasia using pattern-oriented modelling methods (Grimm *et al*. 2005) and spatiotemporal evidence from hundreds of radiocarbon dated fossils and ancient DNA sequences (Fig 1 in S1 Text). Driver-state relationships simulated the effects of climate change and hunting by humans on key ecological processes of extinction: niche lability, dispersal, population growth and Allee effect. Models were coded in Program R (available here: https://doi.org/10.5281/zenodo.5567859) and are described in detail in the Methods in S1 Text.

**Fig. 1.**
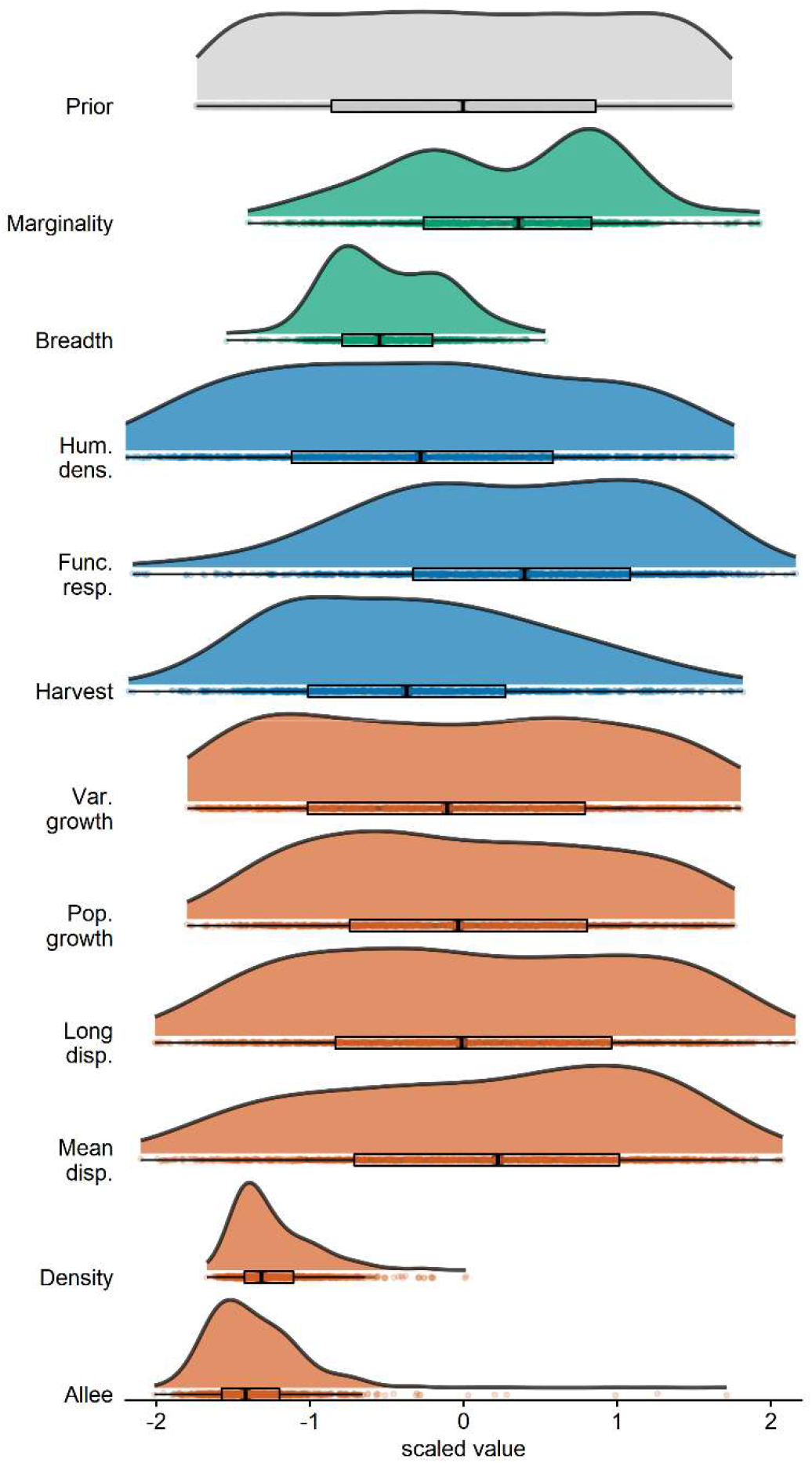
Posterior distribution of parameter values. Distributions for prior values (grey), and posterior values for ecological niche requirements (green), exploitation parameters (blue) and demographic parameters (orange). Values on the y-axis have been scaled so that the maximum height of each density plot is 1. Values on the x-axis are centred and scaled. Box plots show the 25th, 50th and 75th percentiles. Whiskers show maximum values. Ecological niche requirements are distance between average climatic conditions of the occupied niche and the average climatic condition in the study region (Marginality); and breadth of climatic conditions the species can occupy (Breadth). Harvest parameters are percentage of the population that is harvested (Harvest); extent to which harvest follows a Type II to Type III functional response (Func. resp.); and maximum human abundance in a grid cell (Hum. dens.). Demographic processes are Allee effect, maximum abundance (Density); maximum dispersal distance (Long disp.); mean dispersal rate (Mean disp.); maximum population growth rate (Pop growth); and variation in population growth rate (Var growth). See Methods in S1 Text and Appendix 3 in Fordham and Brown (2020) for more details.

### Woolly mammoth niche

Radiocarbon dated and georeferenced fossils for the woolly mammoth (*M. primigenius*) during the Late Pleistocene and Holocene were sourced from publicly accessible databases and published literature (Fordham & Brown 2020). Their age reliability was assessed (Barnosky & Lindsey 2010) and all reliable ages were calibrated using OxCal (Bronk Ramsey 2009) and the IntCal13 calibration curve (Reimer *et al*. 2013). See Methods in S1 Text.

The TraCE-21 simulation of the transient climate of the last 21,000 years was used to generate monthly mean climatic parameters from 21 ka BP to 0 BP (Fordham *et al*. 2017). HadCM3 paleoclimate simulations from 60 to 21 ka BP (Singarayer & Valdes 2010) were harmonized with TraCE-21 simulations and resampled to a 1 × 1° resolution (Methods in S1 Text). We intersected fossil locations and time periods (calibrated age ± 1 SD) with paleoclimate simulations of climatic parameters that affect the population dynamics of large herbivores in polar regions (Sæther 1997) and characterized an n-dimensional hypervolume of climatic suitability through time (Fig 2 in S1 Text), generating a biologically relevant representation of the climate history over which woolly mammoths were present at fossil site (Nogués-Bravo 2009). The resulting hypervolume of climate suitability, which approximates the fundamental niche of the woolly mammoth (Nogués-Bravo 2009), was exhaustively subsampled (Methods in S1 Text). This allowed the realized niche of the woolly mammoth (Fordham *et al*. 2016a) to be identified using process-explicit macroecological modelling (see below). Subsampling was done with Outlier Mean Index (OMI) analysis, using plausible bounds of climatic specialization and niche breadth (Dolédec *et al*. 2000).

**Fig 2.**
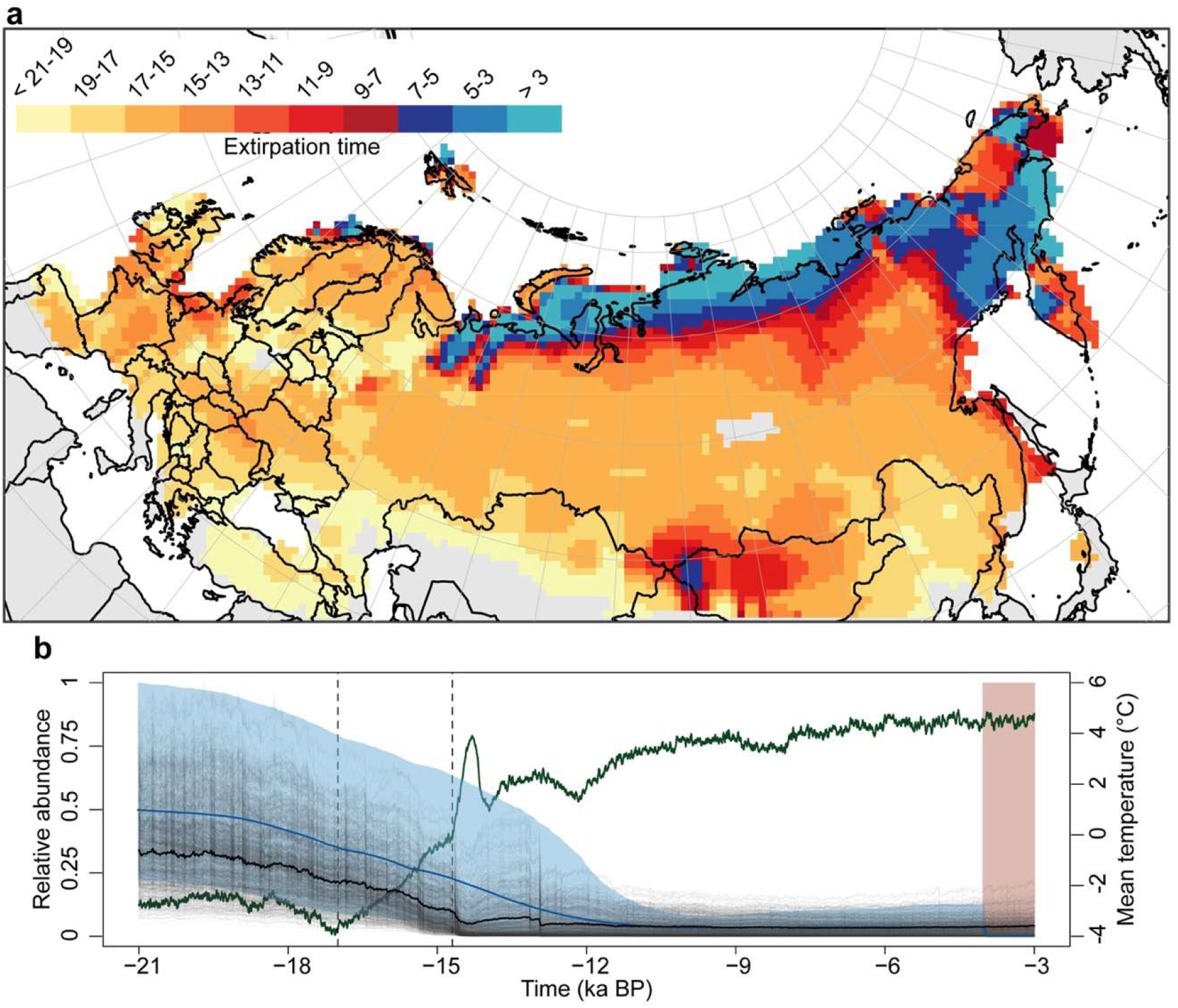
Population decline and extinction of the woolly mammoth in Eurasia. (**A**) Simulated timing of extirpation (model weighted) for woolly mammoth in Eurasia (ka BP). **(B**) Ancient DNA estimates of effective population size (N_e_) (blue line = mean, blue banding = 95% CI) and simulated total population size (black line = weighted mean, grey lines = range) rescaled between 0 and 1. Green line shows change in mean annual temperature for Eurasia. Vertical dashed lines show Heinrich 1 and 14.7 k climatic events. Red band shows estimated time of mammoth extinction.

Spatial projections of climate suitability for woolly mammoths in Eurasia were generated from 21 ka to 0 BP at 25-year time steps for each realized niche (n = 1,862 subsamples of the full hypervolume of climate suitability), using a standard maximum entropy method (Phillips *et al*. 2006), accounting for latitudinal variation in grid-cell size and temporal variation in the proportion of a cell that are land or sea ice (Methods in S1 Text).

### Humans

The peopling of Eurasia by Palaeolithic humans, and their relative abundance, was modelled using a Climate Informed Population Genetics Model (CISGEM) that accurately reconstructs arrival times of anatomically modern humans and current-day distributions of global and regional genetic diversity (Eriksson *et al*. 2012; Raghavan *et al*. 2015). CISGEM simulates local effective population sizes (N_e_) as a function of genetic history, local demography, paleoclimate and vegetation. Like other numerical models of early human migration (Timmermann & Friedrich 2016), arrival, occupancy and density (here N_e_) are forced by spatiotemporal estimates of climate and sea level changes over the past 125 k years. To account for parameter uncertainty in spatiotemporal projections of N_e_ we ran thousands of different models, each with a unique combination of parameter settings (based on established upper and lower confidence bounds; Eriksson *et al*. 2012), selected using Latin hypercube sampling (McKay *et al*. 1979). These N_e_ values were scaled between 0 and 1 and used in the process-explicit macroecological model as potential spatiotemporal measures of relative abundance (n = 10,000 potential scenarios of migration and population growth). CISGEM and its application is described in detail in the Methods in S1 Text.

### Climate-human-woolly mammoth interactions

Extinction and colonization dynamics were simulated as landscape-level population processes, operating at 25-year generational time steps from 21 ka BP. Models centred on ‘best estimates’ for demographic processes (population growth rate and its variance, dispersal, Allee effect), environmental attributes (niche breadth and climatic specialisation) and threats (human abundance and rates of exploitation) were varied across wide but plausible ranges using Latin hypercube sampling of uniform probability distributions, to provide a robust coverage of multi-dimensional parameter space (Fordham *et al*. 2016b). This procedure produced > 90,000 conceivable models (parameterizations) with different combinations of rates of climate change, hunting by humans and demographic processes. Each model was run for a single replicate (Prowse *et al*. 2016). The structure and parameters of the process-explicit model are described in detail in the Methods in S1 Text.

Pattern-oriented modelling (Grimm *et al*. 2005) was used to evaluate different model versions and parameterization, by cross-matching simulations with inferences from paleo-archives (Fordham *et al*. 2020), using Approximate Bayesian Computation (ABC; Csilléry *et al*. 2010). Specifically, simulations of range and extinction dynamics were validated against a four-parameter multivariate target, consisting of trend in total population size (based on N_e_), inferred from ancient DNA (Lorenzen *et al*. 2011); time and location of range-wide extinction, inferred from the fossil record (corrected for the Signor-Lipps effect; Saltré *et al*. 2015); and occupancy at fossil sites. The top 1% of feasible parameterizations of climate-human-woolly mammoth interactions were retained (n = 900) and used to generate ensemble averaged estimates of spatial abundance, timing of extirpation (extinction at the grid cell), total population size, probability of occupancy and hunting rates. Estimates were weighted by the Euclidean distance of the model from the idealized targets.

Our process-explicit modelling approach also permits possible alternatives to past events to be simulated and the biological consequences assessed (Fordham *et al*. 2020). For example, we held human-hunting parameters constant at zero harvesting in the best 1% of validated models and then analysed the effect of this constraint on dynamical processes and emergent patterns, and compared these to model simulations of climate and human interactions with the woolly mammoth.

### Statistical analysis

To identify and evaluate the processes and drivers that caused the initial population collapse of woolly mammoth, and later the susceptibility of small residual populations to eventual extinction (declining and small population paradigms; Caughley 1994), we discretised results from the best 900 models into three distinct climatic periods (T1 = 21-15 ka BP, T2 = 15-11 ka BP and T3 = 11-5 ka BP) (Clark *et al*. 2012) and three sub-regions (Europe, Asia, Beringia). For each period and region (including all of Eurasia) we computed the magnitude and pace of climatic change, human population growth and expansion (Fordham & Brown 2020). We calculated Expected Minimum Abundance (EMA) at the termination of each period (T1-T3) for each region. EMA quantifies risks of both overall population decline (quasi-extinction) and total extinction (McCarthy & Thompson 2001). Statistical learning models (Wright & Ziegler 2017) were used to identify spatiotemporal determinants of extinction risk (Pearson *et al*. 2014). Spatiotemporal autocorrelation between climatic and human drivers of extinction was calculated using the Lee’s L Statistic (Lee 2001). Phase synchrony and peak coincidence was calculated using the synchrony package for R (Gouhier & Guichard 2014).

## Results

Successfully simulating vital aspects of the range collapse and extinction of woolly mammoth, inferred from paleo-archives, required highly constrained environmental attributes (climatic requirements) and demographic mechanisms in our process-explicit model (Fig 1). This indicates that ecological niche requirements and individual fitness at small population size are likely to have been crucial drivers of the extinction dynamics of woolly mammoth in continental Eurasia. Our prior-to-posterior checks show that non-marginal niches (niches with low specialisation), low maximum abundances and a small Allee effect is needed to simulate both the timing and rate of population decline (based on ancient DNA) and the location and timing of range-wide extinction (based on the fossil record) (Table 1 in S1 Text). By comparison, posterior distributions for exploitation parameters (including harvest rate) more closely matched prior distributions (Fig 1).

The best models (top 1% of all models, which most closely matched the validation targets) reconstructed a north-eastward range contraction for woolly mammoth from 19 ka BP, with extirpation in most of Europe by ∼ 14 ka BP (Fig 2), with the exception of refugial areas in what is now Britain, northern France and Belgium (as well as pockets in the Netherlands and Denmark), where patches of steppe-tundra ecosystems are likely to have persisted in favourable climatic areas until the termination of the Pleistocene (Binney *et al*. 2017). These best models simulated an accelerated rate of range collapse in Asia following rapid warming at ∼ 15 ka BP (S1 Movie), with populations persisting within the Siberian refugia of Taimyr, Beringia and the Yamal Peninsula, in accordance with fossil remains (Stuart *et al*. 2004; Stuart 2005). Furthermore, these models accurately projected a steep decline in total population size during the Late Pleistocene, as inferred here from ancient DNA estimates of effective population size (Fig 2). Our simulations, however, did not project a continent-wide extirpation during the early Holocene.

To converge on the validation targets, simulations needed woolly mammoth to persist in Beringia, along the Kara Sea (including the Taimyr Peninsula) and, possibly in northern Fennoscandia (following the retreat of glacial ice sheets), until at least the mid-Holocene (Fig 2). Model agreement for persistence in these mainland Arctic refugia until the mid-Holocene was generally high in the best 1% of models (S1 Movie), pinpointing locations of Holocene-age refugia. These refugial locations are likely, given the incompleteness of the fossil record (Fig 3 in S1 Text) and low numbers of mammoths projected during the Holocene (Fig 2). In contrast, refugia projected in the high elevation plateaux of southern Asia, and in Svalbard, during the Holocene had low model agreement: on average < 9% probability of occurrence (S1 Movie).

**Fig 3.**
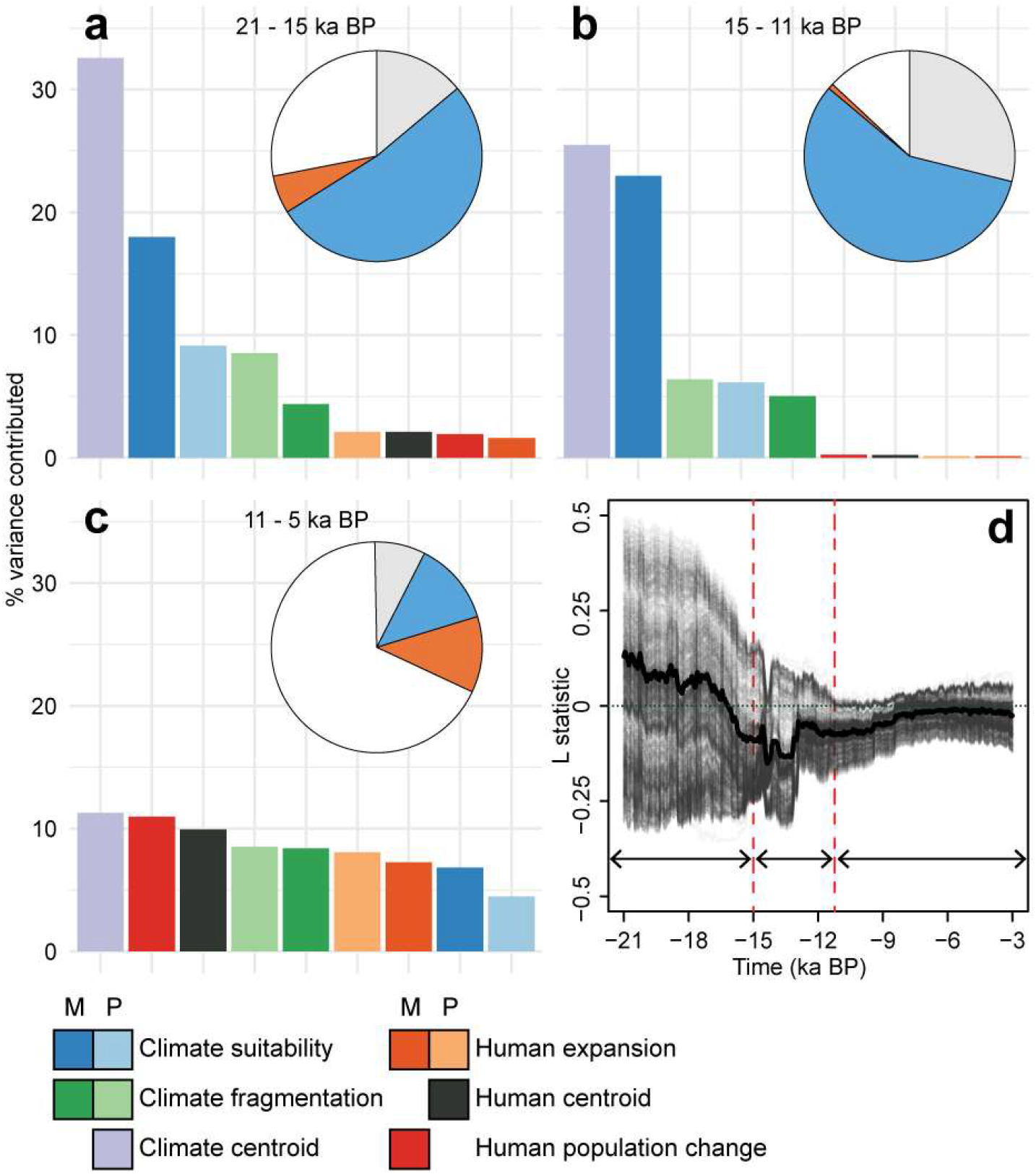
Effects of humans and climate on the decline of woolly mammoth across Eurasia. Drivers of expected minimum for the periods 21-15 ka BP (**A**), < 15-11 ka BP (**B**), < 11-5 ka BP (**C**) for the 1% best validated models. Pie charts show variance explained (%) by climate (blue), humans (orange) and area of occupancy at the start of each period (grey). Histograms show contribution to explained variance for magnitude (M) or pace (P) of variables detailed in Appendix 1 of Fordham and Brown (Fordham & Brown 2020). White areas of pie chart represent unexplained variance. (**D**) shows the relationship (Lee’s L statistic) between human abundance and climate suitability for mammoths across time for Eurasia (black line = weighted mean, grey lines = individual model runs). Positive values indicate positive correlation, whilst negative values indicate a decoupling of the variables (negative correlation). Fig 7 in S1 Text shows variable importance for area of occupancy.

### Spatiotemporal determinants of extinction

Across Eurasia, climatic change during the last deglaciation and the Holocene affected the geography of human and woolly-mammoth abundance differently based on the best models of climate-human-woolly mammoth interactions (Fig 3). Accelerated warming following Heinrich Event 1 (∼ 17.5 ka BP) resulted in climatically preferred conditions (temperature and precipitation) for mammoths and humans becoming decoupled (mean Lee’s L statistic of spatial autocorrelation = – 0.07 between 17.5 and 15 ka BP), due likely to humans responding to environmental change by colonizing and remaining resident in new niches (Giampoudakis *et al*. 2017), and the woolly mammoth retreating to the coldest areas of its climatic niche, where conditions for people were most harsh.

Using statistical learning analysis to identify spatiotemporal determinants of extinction risk from these best models, we show that during the last deglaciation, climatic shifts alone explained 52 and 57 % of the variance in expected minimum abundance (EMA) of woolly mammoth in Eurasia at 15 ka BP and 11 ka BP, respectively (Fig 3). Larger magnitudes of change in the climatic conditions suitable for woolly mammoth persistence, along with a faster pace of loss of these conditions, resulted in increased extinction risk during the periods 21-15 ka BP (T1) and 15-11 ka BP (T2) (Fig 4 in S1 Text). The magnitude and pace of human population growth became as important as climatic change during the Holocene in influencing the EMA of woolly mammoth in Eurasia at 5 ka BP (Fig 3). A lower explained variance in the Holocene compared to earlier time periods occurs because pattern-oriented validation procedures retain models that simulate mechanisms of Late Pleistocene range collapse and population decline (those occurring in T1 and T2) having a long-lasting effect on the timing and location of extinction in the Holocene (T3; 11-5 ka BP).

**Fig 4.**
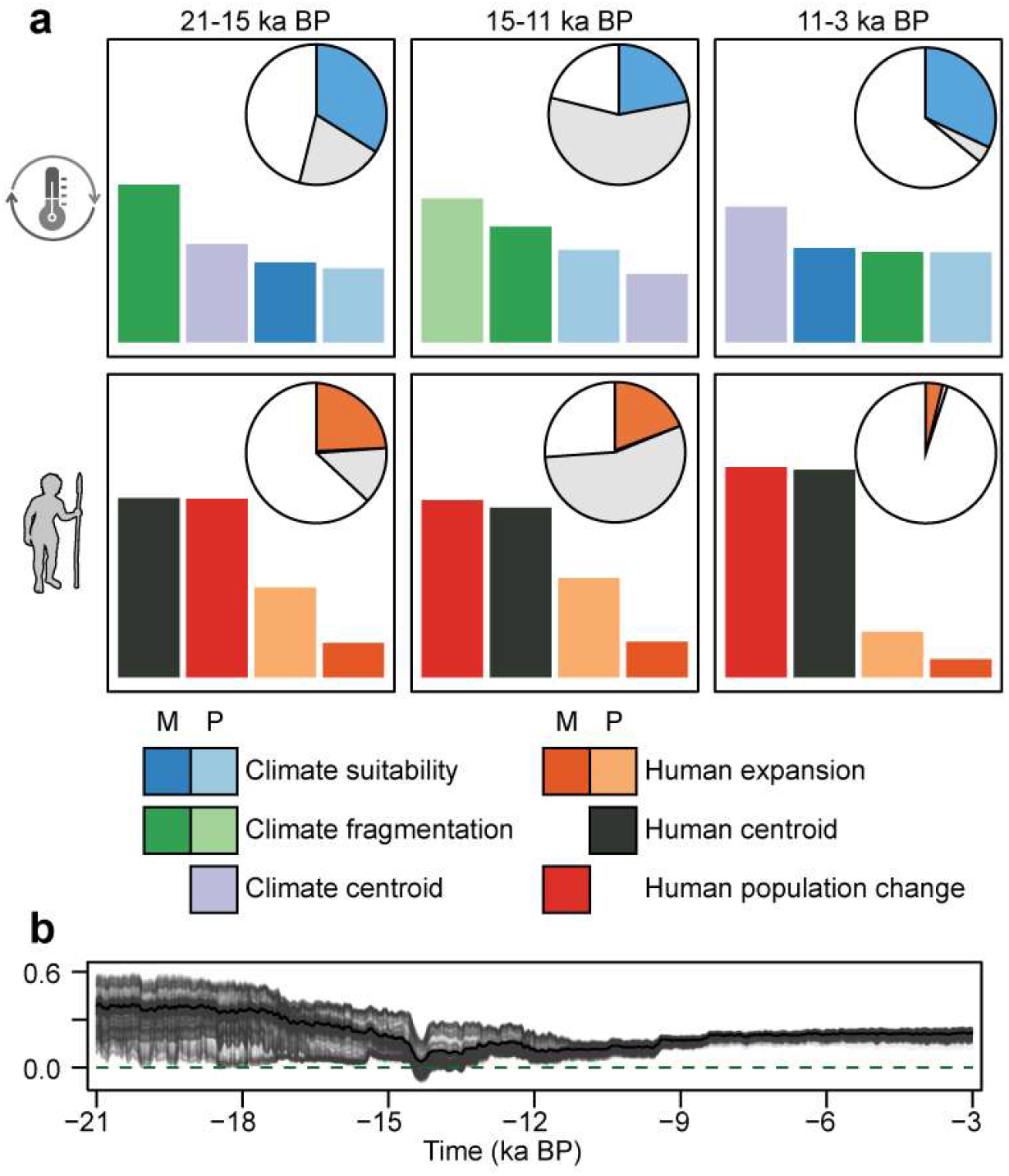
Extinction risk in Europe for woolly mammoth due to humans and climate. (**A**) Variable importance (histograms) and variance explained (pie charts) for climate (top row) and human parameters (middle row). Fig 2 describes the legend. (**B**) Lee’s L statistic of autocorrelation between human abundance and climate suitability for mammoths.

Demographic responses of woolly mammoths to impacts of humans and climate were spatiotemporally heterogenous, with these differences being essential for setting the time and place for extinction. Populations of woolly mammoths in distinct regions of Eurasia experienced very different magnitudes of climatic and human impacts over time (Fig 5 in S1 Text), suggesting that the dynamics of extinction determinants were labile. Accurately reconstructing inferences of range collapse and population change from fossils and ancient DNA required that humans impacted woolly mammoth prior to the Holocene in Europe.

**Fig 5.**
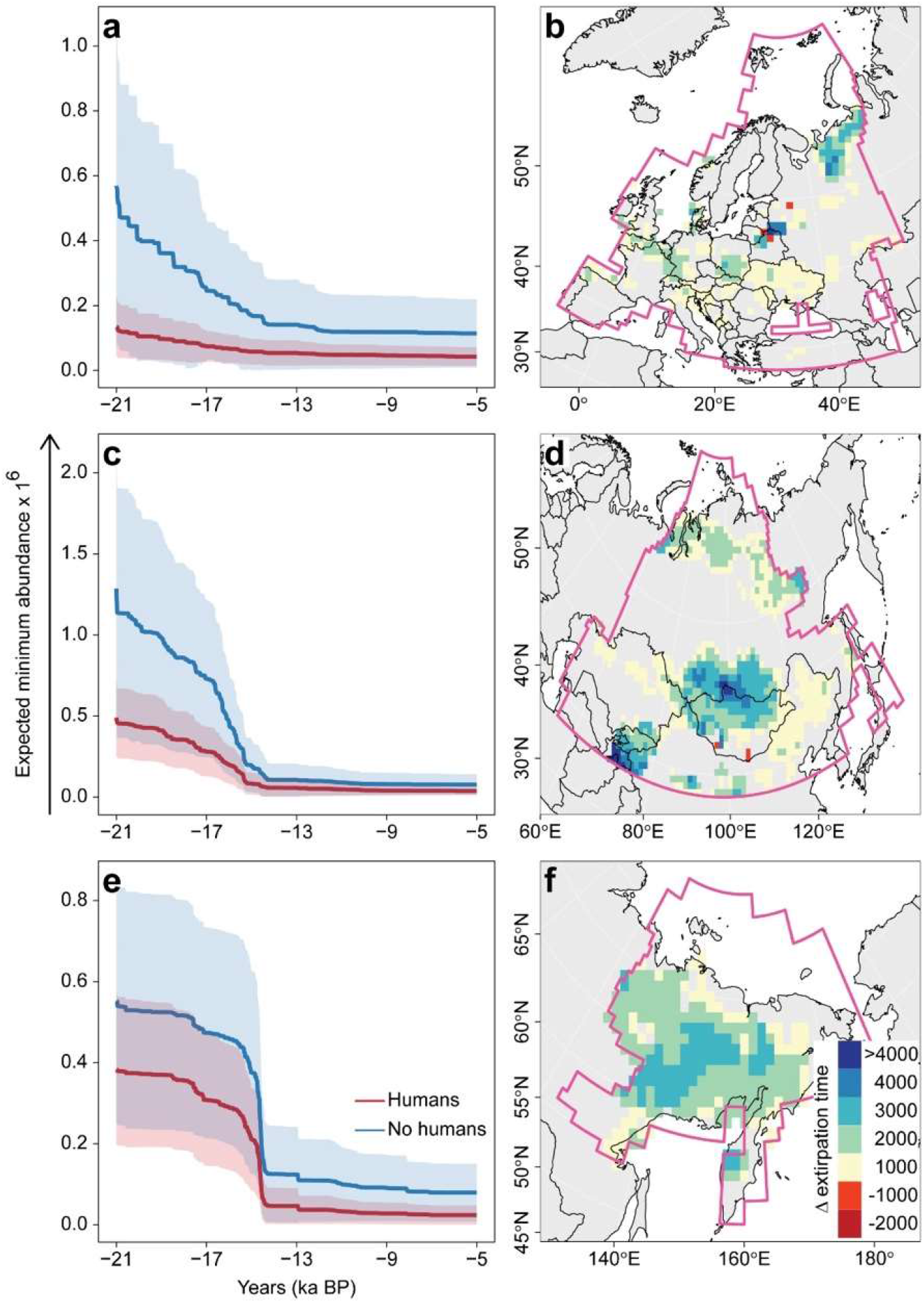
Footprint of humans on the extinction dynamics of woolly mammoths. Relative change in expected minimum abundance of woolly mammoth in response to climate change and exploitation from 21 ka BP (Humans) and a counterfactual no-exploitation scenario (No humans) for Europe (**A**), Asia (**C**) and Beringia (**E**). Maps show the difference in timing of extirpation for woolly mammoths in the absence of hunting by humans (i.e. No humans – Humans) for Europe (**B**), Asia (**D**) and Beringia (**F**). Positive values indicate a later extirpation date in the absence of humans. Differences of ± 500 years are not shown. Pink lines mark regional boundaries.

The magnitude and pace of climatic and human impacts during the last deglaciation in Europe explained relatively similar amounts of variance in EMA of woolly mammoth (Fig 4). During this period, humans and mammoths responded differently to climatic change in Europe (mean Lee’s L = 0.21 for the last deglaciation, indicating low autocorrelation), confirming direct and important roles of hunting and climatic change on mammoth EMA. Together these results indicate that humans had a sustained effect on the population dynamics and extirpation of woolly mammoths in Europe. For the few populations in northern Fennoscandia that were simulated to have colonized the region following the retreat of glacial ice sheets (S2 Movie), gradual climatic change during the Holocene was the greatest threat to simulated persistence (Fig 4) not hunting by humans, which remained relatively low (S3 Movie), owing to small population densities of humans in northern Fennoscandia (S4 Movie).

In Asia and Beringia, the role of humans on the range collapse and extirpation of woolly mammoth more closely mirrored the pattern of continent-wide extinction observed for Eurasia, with humans having a proportionately larger threatening influence on EMA by the Holocene (Fig 5 in S1 Text). In Beringia, climatic changes during the last deglaciation and Holocene affected humans and mammoths more similarly than for other regions of Eurasia (Fig 5 in S1 Text; mean Lee’s L = 0.44 for the last deglaciation; and 0.35 for the Holocene), indicating greater synchrony in shifts in range and abundance.

### Climate-human interactions

Counterfactual models with no mammoth exploitation by humans provide further evidence that humans acted to hasten the timing of range collapse and extinction of woolly mammoths. We show that in the absence of humans and their interactions with climatic change, woolly mammoths would have been more abundant across time (including during the Late Pleistocene), and populations would have persisted for much longer, perhaps even avoiding extinction within climatic refugia (Fig 5). Mid-to-late Holocene population sizes of woolly mammoths were much larger in simulations in the absence of human harvesting (Table 2 in S1 Text), causing a 24% increase in persistence beyond 3.8 ka BP; the estimated time of extinction (95% confidence interval = 4089 to 3450 BP) based on the fossil record (Methods in S1 Text). Although some model simulations with humans on the landscape also did not result in range-wide extinction by 3.8 ka BP (Fig 6 in S1 Text), confidence intervals for EMA intersect zero (Table 2 in S1 Text).

While simulations of mammoth abundance, with and without humans, were asynchronous (except for Beringia prior to 15 ka BP), the extreme peaks and troughs in abundance generally occurred at similar times in the simulations (Table 3 in S1 Text). This was not the case for Europe during the early part of the last deglaciation (21 – 15 ka BP) and Asia during the latter part of the last deglaciation (15 – 11 ka BP), confirming that the strength of the regulatory role of humans on the extinction dynamics of woolly mammoth was regionally and temporally variable.

## Discussion

Macroecological models of mechanistic interactions of climate and humans with the woolly mammoth show that its pathway to extinction was long and lasting, starting many millennia before the final extinction event with successive regional extirpations rather than a rapid range-wide collapse. We show that reconciling spatiotemporal evidence of the decline and extinction of woolly mammoths from fossils and ancient DNA requires quite tight demographic and niche constraints, and spatiotemporally variant rates of climatic change and human impacts (and their synergy) since the Last Glacial Maximum. Our simulations reveal that human population growth and northward migrations during the Late Pleistocene led to the premature extirpation of populations of woolly mammoth in areas of Eurasia that were climatically suitable into the Holocene, hastening climate-driven declines by up to 4,000 years in some regions. They also show that simulating important aspects of the extinction dynamics of woolly mammoths as inferred from paleo-archives requires persistence in mainland Arctic refugia until the mid-Holocene. Thus process-explicit models that continuously simulate the causes of early-stage population declines, and later the susceptibility of small residual populations to eventual extinction, provide new opportunities to unravel ecological mechanisms of extinction that occurred in the ancient past.

The role of humans as a causative agent in the extinction of woolly mammoths is likely to have been both direct and indirect. In addition to exploitation-driven changes in demographic processes, the climate-change-facilitated co-occupation of steppe and forb habitats by humans and woolly mammoths could have affected metapopulation structures and processes by interrupting sub-population connectivity, affecting movement between resource-rich zones (Cooper *et al*. 2015). Posterior distributions for exploitation parameters (including harvest rate) for Eurasia-wide simulations more closely matched prior distributions when compared to some ecological niche and demographic parameters, indicating reduced parameter importance. The reduced importance of the role of human hunting in the extinction dynamics of woolly mammoth at a continental scale likely reflects variable rates of hunting of woolly mammoths by humans in space and time and a likelihood that, to some extent, humans impacted mammoth metapopulation processes independent of hunting (e.g., via habitat change, with hunting being most damaging at specific ‘pinch-points’ such as movement corridors).

Correctly simulating inferences of extinction dynamics from fossils and ancient DNA, required the survival of woolly mammoth in mainland Arctic refugia until the mid-Holocene, some 5,000 years longer than previously thought based on fossil evidence alone (Stuart *et al*. 2004; Nikolskiy *et al*. 2011; Dehasque *et al*. 2021). Extirpation and extinction events for megafauna are commonly revised as younger fossils and environmental DNA are discovered, often causing persistence to be extended by several millennia (Haile *et al*. 2009; Wang *et al*. submitted). This is because actual extinction dates are often underestimated using date of last fossil appearance, which usually records the last time a species was abundant not last occurrence (Mann *et al*. 2019). This raises the real possibility that populations of woolly mammoth persisted in mainland arctic refugia until the mid-Holocene, as predicted by our model. Especially given that population abundances of mammoths during the Holocene would have been low (Fig 2), indicator species of mammoth persisted in Siberia during the mid-Holocene (Boeskorov 2020), the refugial areas that we pinpoint remain poorly sampled (Fig 2 in S1 Text), and environmental DNA evidence of woolly mammoths in Beringia and the Taimyr Peninsula to the mid-Holocene (Wang *et al*. submitted). The recent discovery of persistence of woolly mammoths in Siberia to 3.9 ka using environmental metagenomics (Wang *et al*. submitted) provides an important independent validation of our process-based model (Grimm *et al*. 2005), indicating a strong ability of the simulations to detect hidden refugia and unveil spatiotemporal pathways to extinction.

Although equifinality was avoided in our analyses, using multiple validation targets, our finding that mammoths likely persisted in mainland refugia in Eurasia until the mid-Holocene was dependent on currently available inferences of extinction from the fossil record, estimates of population decline from ancient DNA, and projections of spatially and temporally variant rates of climatic change and human impacts. While the fossil record for woolly mammoth, during the Pleistocene-Holocene transition, is more complete then for many other megafauna species, there are still vast areas of Eurasia which remain poorly represented by fossil samples (Fig 3 in S1 Text). Likewise, projections of population growth and migrations of people during the Late Pleistocene and Holocene do not yet account for topographical processes and cultural changes (Eriksson *et al*. 2012; Timmermann & Friedrich 2016), which could potentially influence our results. Therefore, uncertainty in our estimates of the extinction dynamics of woolly mammoths in space and time is likely to be reduced through new fossil discoveries, and higher spatial resolution projections of the peopling of Eurasia.

The prior and posterior parameter distributions of some demographic and exploitation parameters in the process-explicit models were similar, indicating that a wide range of values for these parameters will reconstruct spatiotemporal evidence of the decline and extinction of woolly mammoths. For these non-identifiable parameters, it is important to consider whether the empirical targets used for pattern-oriented modelling best fit the study animal and system (Gelman *et al*. 2013), which they do for the woolly mammoth. Nevertheless, it does mean that a variety of different parameter combinations can give a close fit to inferences of extinction dynamics from the fossil record and ancient DNA, potentially resulting in different ecological interpretations.

Here we opened a window into late Quaternary biodiversity dynamics using process-based macroecological models, in order to synthesize disparate evidence from paleo-archives (*sensu* Fordham et al. 2020), establishing that ecological pathways to extinction can start many millennia before populations are at critically small thresholds. This reinforces the need for long-term perspectives when testing theories and making generalisations regarding the spatial dynamics of range collapses and extinctions of species. Our results emphasise that extinctions can only be explained by combining the declining and small population paradigms (Caughley 1994). They also highlight that range shifts during the late Quaternary offer distinct opportunities to test the circumstances under which geographic ranges collapse first along the periphery versus those which start within the range interior (Channell & Lomolino 2000), particularly if they are reconstructed using process-explicit models and pattern-oriented approaches (Fordham *et al*. 2021).

Our analyses strengthen and better resolves the case for human impacts as a crucial and chronic driver of early-stage population declines of megafauna, revealing an essential role of humans in population declines of mammoths in Eurasia during the Late Pleistocene; a period when climatic conditions warmed rapidly. In doing so, it refutes a prevalent theory that the role of humans in the extinction dynamics of woolly mammoths was limited to a mid-Holocene coup de grâce (Nogués-Bravo *et al*. 2008), and highlights the importance of disentangling demographic responses to varying biotic and abiotic stressors for metapopulations at regional scales, particularly when assessing species survival under climate change.

## Acknowledgments

F. Saltré, K. Giampoudakis, A. Flórez-Rodríguez assisted with collating and analysing the fossil record and ancient DNA.

## Supporting Information

**S1 Text**. Supporting methods, figures, and tables

**S1 Movie**. Probability of occupancy for woolly mammoth in Eurasia from 21,000 BP

**S2 Movie**. Density estimates of woolly mammoth in Eurasia from 21,000 BP

**S3 Movie**. Harvest rates of woolly mammoth in Eurasia from 21,000 BP

**S4 Movie**. Effective population size of humans in Eurasia from 21,000 BP

